# Phylochemical mapping of natural products onto the plant tree of life using text mining and large language models

**DOI:** 10.1101/2024.02.16.580694

**Authors:** Lucas Busta, Drew Hall, Braidon Johnson, Madelyn Schaut, Caroline M. Hanson, Anika Gupta, Megan Gundrum, Yuer Wang, Hiroshi A. Maeda

**Author notes:** **Corresponding Authors:** Lucas Busta and Hiroshi A. Maeda.

## Abstract

Plants produce a staggering array of chemicals that are the basis for organismal function and diversity and also provide essential human nutrients and medicine. However, it is poorly defined how these compounds have evolved and are distributed across the diverse lineages of the plant kingdom, hindering a systematic view and understanding of plant chemical diversity. Recent advances in plant genome/transcriptome sequencing have provided a well-defined molecular phylogeny of plants, on which the presence of diverse natural products can be mapped to systematically determine their phylogenetic distribution. Here, we built a proof-of-concept workflow via which previously reported diverse tyrosine-derived plant natural products were mapped on to the plant tree of life. Plant chemical-species associations were mined from literature, filtered, evaluated through manual inspection of over 2,500 scientific articles, and mapped onto the plant phylogeny. The resulting ‘phylochemical’ map confirmed several highly lineage-specific compound class distributions, such as betalain pigments and Amaryllidaceae alkaloids. The map also highlighted several lineages enriched in dopamine-derived compounds, including the orders Caryophyllales, Liliales, and Fabales. Additionally, the application of large language models using our manually curated data as a ground truth set showed that post-mining manual processing steps can largely be automated with a low false positive rate. Our study demonstrates that a workflow combining text mining with language model-based processing can generate broader phylochemical maps, which will serve as a critical community resource to uncover key evolutionary events that underlie plant chemical diversity and enable system-level views of nature’s millions of years of chemical experimentation.

**Significance Statement:** This study introduces a novel workflow to map the distribution of natural products onto a phylogeny. It also demonstrates the effectiveness of combining text mining of the literature with large language model evaluation for constructing a ‘phylochemical’ database that can organize vast portions of the chemical diversity present across the plant tree of life.

## Introduction

Plants synthesize a vast array of chemical compounds that are essential for their own physiological processes and adaptation to the ecological niches they inhabit. These plant natural products also serve as vital nutrients, flavors, antioxidants, and neurotransmitters for other organisms. To date, researchers have identified over 200,000 such compounds, with estimates suggesting the existence of more than a million distinct chemicals in the more than three hundred thousand species that make up the plant kingdom.^1,2^ Importantly, about one third of all pharmaceuticals approved in the last 35 years are obtained from plant-derived substances, either directly or as synthetic derivatives inspired by them.^3,4^ Furthermore, roughly two-thirds of the global population depends on plants as primary medicinal resources.^5,6^ Thus, while synthetic combinatorial chemistry has produced extensive chemical libraries for drug discovery, the intricate and specialized nature of plant-based compounds, honed by evolutionary processes, render them a treasured source for new pharmaceutical developments^7–9^ and underscore the need for expanded exploration of natural products across the plant tree of life.

Plant natural products are largely produced by complex networks of metabolic pathways that evolve via various mechanisms including gene duplication, enzyme neofunctionalization, and altered gene expression.^10–13^ Many of these plant pathways and their products occur in lineage-specific patterns and thus are referred to as lineage-specific or specialized metabolism, as compared to primary metabolism which is conserved across most organisms. The taxonomic distributions of plant chemicals have been an active area of study for decades, with contributions coming from diverse research areas including chemotaxonomy, natural products research, botany, drug discovery, and many others. Thus, though the distribution of plant chemicals is complex, a large amount of information is available about the detection of various chemicals in different plant species. However, this wealth of information was collected by many different investigators using diverse approaches (including chromatography, mass spectrometry, nuclear magnetic resonance, infrared spectroscopy, etcetera), and is distributed across a large number of publications.

Recent studies integrating molecular phylogenies with metabolite analyses have showcased the value of exploring chemical distributions across the plant tree of life.^14–16^ These analyses have focused on either a broad range of species with a targeted set of metabolites, or a broad range of metabolites across a specific group of species. Such targeted analyses have, for example, helped predict chemical diversity hotspots,^17^ helped us understand some of the genetic and evolutionary basis for specialized metabolism,^18,19^ and uncovered variant metabolisms with industrial or agricultural applications.^20^ These outcomes alone are exciting but also hint at potentially transformative outcomes that could result from broader analyses of plant chemical distributions across the tree of life, i.e., by incorporating wide ranges of both metabolites and species. The ingredients for such untargeted analyses are already available in the form of (i) massive molecular phylogenies enabled by recent advances in sequencing technology and (ii) decades of literature documenting the presence of lineage-specific plant chemicals. Unfortunately, a bottleneck exists in the highly time consuming and laborious tasks of systematically extracting compound-species associations from the literature and documenting them in a way that enables them to be mapped onto a phylogeny. By overcoming this bottleneck, we can gain a much more holistic perspective of plant chemical diversity that will help us (i) understand the evolutionary history of diverse natural products and underlying metabolic innovations, (ii) conduct targeted searches for new bioactivities, and (iii) derive novel hypotheses and design experiments aimed at understanding the physiological and ecological functions of plant specialized metabolites.

Several other groups have worked toward systematically extracting compound-species associations from literature. These efforts have led to several natural product databases, including Duke’s Plant Natural Product, KNAPSAcK, the Dictionary of Natural Products, and NAPRALERT.^2,21,22^ These databases are excellent tools for targeted searches and provide detailed information about bioactivities, mass spectral signatures, physical properties of chemicals, and in some cases compound-species associations. Nonetheless, these resources are designed for targeted searches for a specific compound(s) or a specie(s) of interest and not for untargeted analyses that require comprehensive phylogenetic mapping of plant compounds. To bridge this gap, we have developed a novel workflow (**Figure 1**) for creating a ‘phylochemical’ database in which chemical diversity is organized in the context of a plant phylogeny and all data are simultaneously accessible. This workflow consists of (i) text mining to identify potential associations between chemical compounds and specific plant species, (ii) evaluation of those associations, and (iii) mapping of positive hits on molecular phylogeny to populate a phylochemical database (**Figure 1**). Here, we first used text mining and manual curation to identify and evaluate over 3,000 putative associations between specific plant species and specific tyrosine-derived compounds. We then analyzed the distribution of the tyrosine-derived compounds across various plant lineages to gain insights into their specialized metabolic pathways. Finally, we tested the potential of large language models to complement our text mining methods, revealing their effectiveness in processing and accurately categorizing candidate compound-species associations. Overall, our study demonstrates that a workflow combining text mining with language model-based processing can open the bottleneck preventing the construction of broader phylochemical maps that span large portions of both chemical and plant diversity.

**Figure 1.**
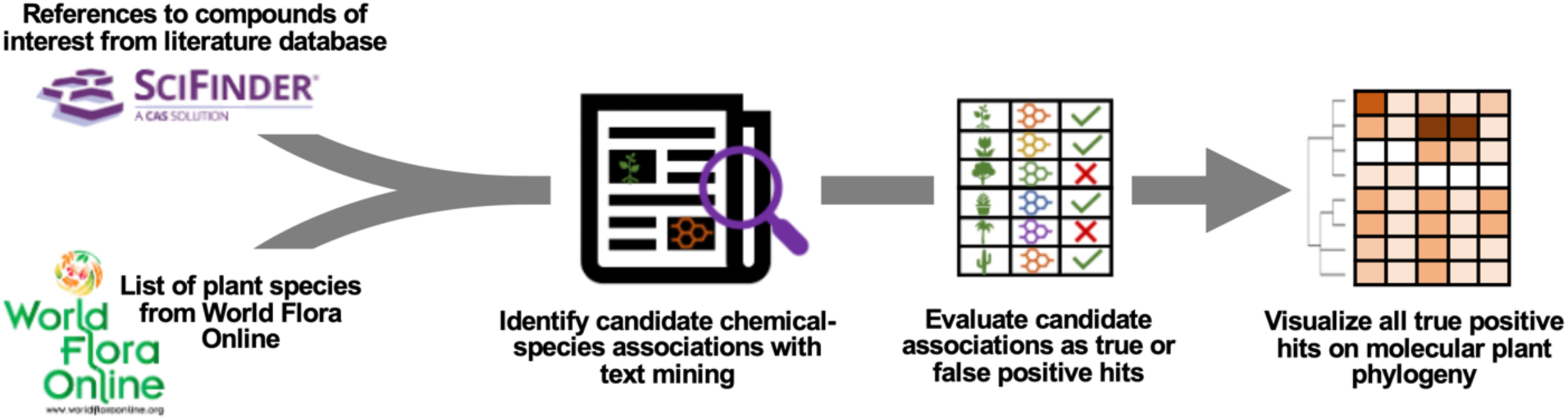
Overview of the workflow for identifying compound-species associations. From left to right, the workflow consists of gathering relevant articles from a literature database together with a list of accepted plant species names, then applying text mining to identify co-occurrences of the names of chemical compounds and chemical species in the titles or abstracts. Next, the co-occurrences (candidate compound-species associations) are evaluated to isolate only associations that are supported with experimental evidence. Finally, the supported compound-species associations are mapped on to a phylogeny.

## Results

### Text mining and manual curation to identify compound-species associations

Our first aim was to identify previously reported associations between tyrosine-derived compounds and specific plant species. Tyrosine-derived metabolites were useful in developing a phylochemical mapping workflow because some tyrosine-derived compounds are broadly distributed while others are highly lineage-specific.^23^ The latter include betalains and Amaryllidaceae alkaloids, which serve as internal positive controls. We began by compiling a list of 71 tyrosine-derived plant natural products, based on prior literature and the KEGG plant chemical compound list,^23,24^ and grouped them based on their upstream precursor molecules or commonly used compound class names (e.g., ‘dopamine-derived compounds’, ‘Amaryllidaceae alkaloids’, etcetera, **Supplemental Table S1**). We then used the Chemical Abstracts Service (CAS) SciFinder database^25^ to retrieve references, including titles and abstracts, for articles that mentioned the names of the 71 tyrosine-derived natural products. Subsequently, we filtered the SciFinder search results to include only the articles that were associated with the keyword ‘plant’ or ‘taxonomy’ in the SciFinder database, thereby eliminating a large number of clinical reports, which were likely irrelevant to our plant-derived chemical focus. Even after filtering, eight compounds, including epinephrine (adrenaline), noradrenaline, dopamine, and colchicine had over 10,000 hits (**Supplemental Figure S1**), most of which appeared to be due to human clinical studies of these widely used pharmaceuticals. It seemed likely that many of the co-occurrences of these compounds and plant species names in the text of these articles would not reflect endogenous production in plants, so we excluded these compounds from further analysis. The references for the remaining 63 compounds were then filtered for articles that were written in English and had titles or abstracts mentioning at least one plant species name included in The World Flora Online (WFO) Plant List—an open access resource for the most comprehensive plant list maintained by the global community of taxonomy experts (https://wfoplantlist.org/plant-list). Overall, this process resulted in 2,571 pertinent references, for which titles and abstracts were downloaded for further analysis.

In the next phase of our study, we assessed the accessibility of full texts for the 2,571 references using PubChem, hosted by the National Center of Biotechnology Information (NCBI). Unfortunately, full texts were available via NCBI for only approximately 300 articles, with many of the other articles’ full texts being located behind paywalls. Consequently, we first turned our attention to examining the titles and abstracts of the articles for compound-species associations using automated, pattern matching-based text mining. We recorded any occurrence of a taxon name (genus and species names together) within articles that mentioned one of the 63 tyrosine-derived compounds as an indicator of potential presence of the compound in that taxon, yielding 3,631 candidate associations (**Supplemental Table S2**). To integrate these findings with phylogenetic data, we mapped the associations onto a phylogeny of orders within angiosperms.^26^ Branches without reported compounds were pruned for clearer visualization (**Figure 2A**). To account for the varying species counts per order and the disparate number of articles for each compound, we introduced a normalized metric that highlights a compound’s lineage specificity. This metric reflects the total number of reports for a compound per order, divided both by the total number of reports of that compound and by the species count within that order. The resulting phylochemical map showed the expected lineage-specific occurrences of the positive controls (betalains and Amaryllidaceae alkaloids in Caryophyllales and Asparagales orders, respectively), among other patterns (**Figure 2B**). This result demonstrates the ability of text mining to extract compound-species associations and capture their overall phylogenetic distributions.

**Figure 2:**
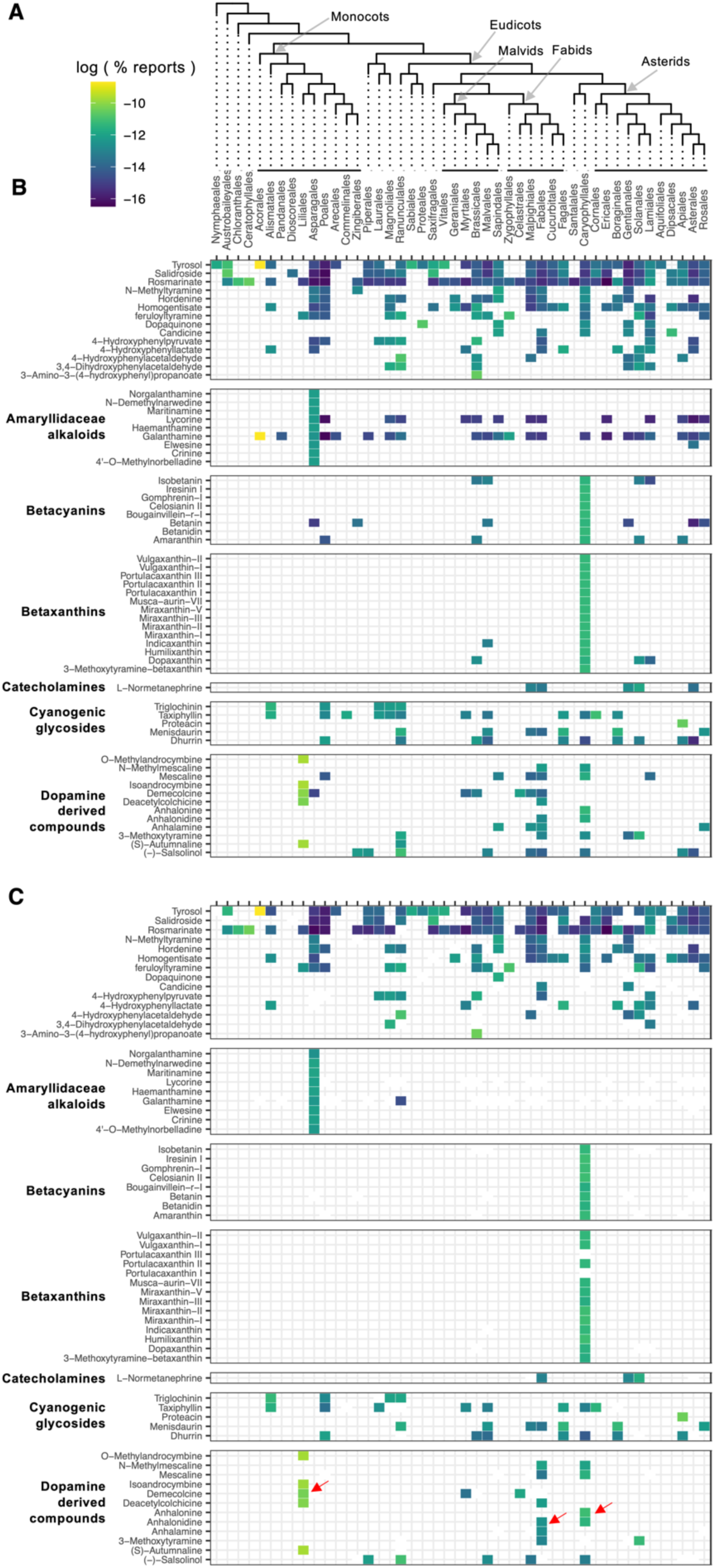
Map of tyrosine-derived compounds across the angiosperm phylogeny. **A.** Cladogram of angiosperm orders. We mapped compound-species associations onto a previously published angiosperm order phylogeny,^26^ removing branches where no compounds were detected for better clarity. **B.** All putative compound-species associations. **C.** All true positives. In B and C, to adjust for differences in species count per order and the number of articles for each compound, we used a normalized metric to color the heat map. This metric is the log of the number of reports for a compound in each order, divided by the total reports of that compound and by the species count in the order, thus emphasizing the compound’s specificity to certain lineages. White boxes in C represent associations that were present in B but were all false positives and thus do not appear in the true positive map.

To assess the reliability of our text mining approach, each of the 3,631 candidate compound-species associations identified were manually reviewed. The review involved manually downloading and searching the full texts, figures, and tables of the 2,571 articles to determine whether each association was a true positive (indicating the compound was indeed present in the species) or a false positive (where the compound and species co-occurred in the text without evidence of the species producing the compound). We considered instances in which the source article reported the pipeline-identified compound-species association either in the abstract, the main text, or in a figure/table as true positives. Our manual evaluation revealed that, of the 3,631 associations detected via text mining, 2,409 were true positives (66.3%, **Figure 2C**), 863 were false positives (23.7%, **Supplemental Figure S2**), and 359 cases (10%) remained ambiguous due to unclear language or indeterminate figures. Overall, the revised map showing only these true positives (**Figure 2C**) contained very similar patterns to the original map (**Figure 2B**), though a number of associations for betalains and Amaryllidaceae alkaloids were eliminated outside of Caryophyllales and Asparagales.

During our manual evaluation of the literature, several patterns emerged that enabled us to distinguish several types of false positive associations. False positives often included: (i) non-original mentions of compounds without supporting data, (ii) discussions of biosynthetic pathways, (iii) mentions of enzyme assays related to a compound without reporting the compound itself, (iv) reports of a compound’s absence in a species, and (v) references to compounds in a context of artificial presence or absence (**Supplemental Table S2**). For example, instances of ‘non-original mention’ were flagged when papers cited a compound without providing new evidence of its presence in the species. Similarly, ‘biosynthetic precursor/derivative’ hits occurred when articles discussed related metabolic pathways or derivatives without confirming the compound’s accumulation. For example, a study mentioned ‘homogentisic acid’ while investigating its metabolic derivatives from *Miliusa balansae* but did not provide evidence of homogentisic acid accumulation.^27^ Some false positives also derived from sources reporting enzymes related to a given compound, such as a large number of papers referencing galantamine, an Amaryllidaceae alkaloid used to treat early-stage Alzheimer’s disease^28^, as a positive control in acetylcholinesterase inhibition assays using the extracts from various other plants.^29,30^ Interestingly, the text mining pipeline also produced false positive hits from papers describing the absence of a compound (for example, papers using targeted metabolomics approaches that reported ‘not detected’). Similarly, broad surveys of multiple species that reported the detection of multiple compounds resulted in false positives. For example, one study surveyed ten Amaryllidaceae species for various compounds including galantamine in *Clivia miniata*^31^; however, galantamine was not detected in *C. miniata* but found in different species presented in the same table. Finally, some false positives resulted from studies using transgenic plants or artificial introduction of compounds to a heterologous species for the purposes of screening enzyme activity or inhibition. These examples and observations will help reduce false positive hits in future iterations of extracting true compound-species associations.

### Exploration of tyrosine-derived compounds across the plant phylogeny

Having compared maps of all putative associations against maps of only true positive hits, we next examined patterns in the true positive map in detail and observed several surprising patterns. Unlike compounds with lineage-specific occurrence such as betalains and Amaryllidaceae alkaloids, we observed that other tyrosine-derived compounds were unexpectedly distributed across multiple lineages, despite them being generally considered as lineage-specific specialized metabolites. Some examples are the tyramine-derived compounds N-methyltyraminemethyl-4-tyramine and hordenine, which were detected in twelve and six different plant orders, respectively, many of which were only distantly related to each other (e.g., Asparagales from monocots and Gentianales and Sapindales from eudicots; **Figure 2**).

To further experimentally validate the somewhat unexpected result described above, we conducted targeted chemical investigations of hordenine and related compounds (**Figure 3A**). We chose plant species that were available in the Botany Greenhouse at UW-Madison and belong to the same lineages reported to have hordenine, harvested live tissues, and analyzed metabolites using liquid chromatography-high-resolution mass spectrometry (LC-HRMS). We found that hordenine and its precursors (i.e., tyramine, N-methyl-4-tyramine) were indeed present in diverse lineages of plants; four orders (Poales, Sapindales, Gentianales, and Caryophyllales) with at least one species in each order testing positive for hordenine production (**Figure 3B**). Some non-Poaceae species accumulated comparable or even higher levels of hordenine and its precursors than *Hordeum vulgare* (Poaceae, **Figure 3C**). These measurements showed that hordenine is not restricted to one plant family and illustrated the ability of the pipeline to provide unexpected, yet accurate phylogenetic distributions of certain chemicals.

**Figure 3.**
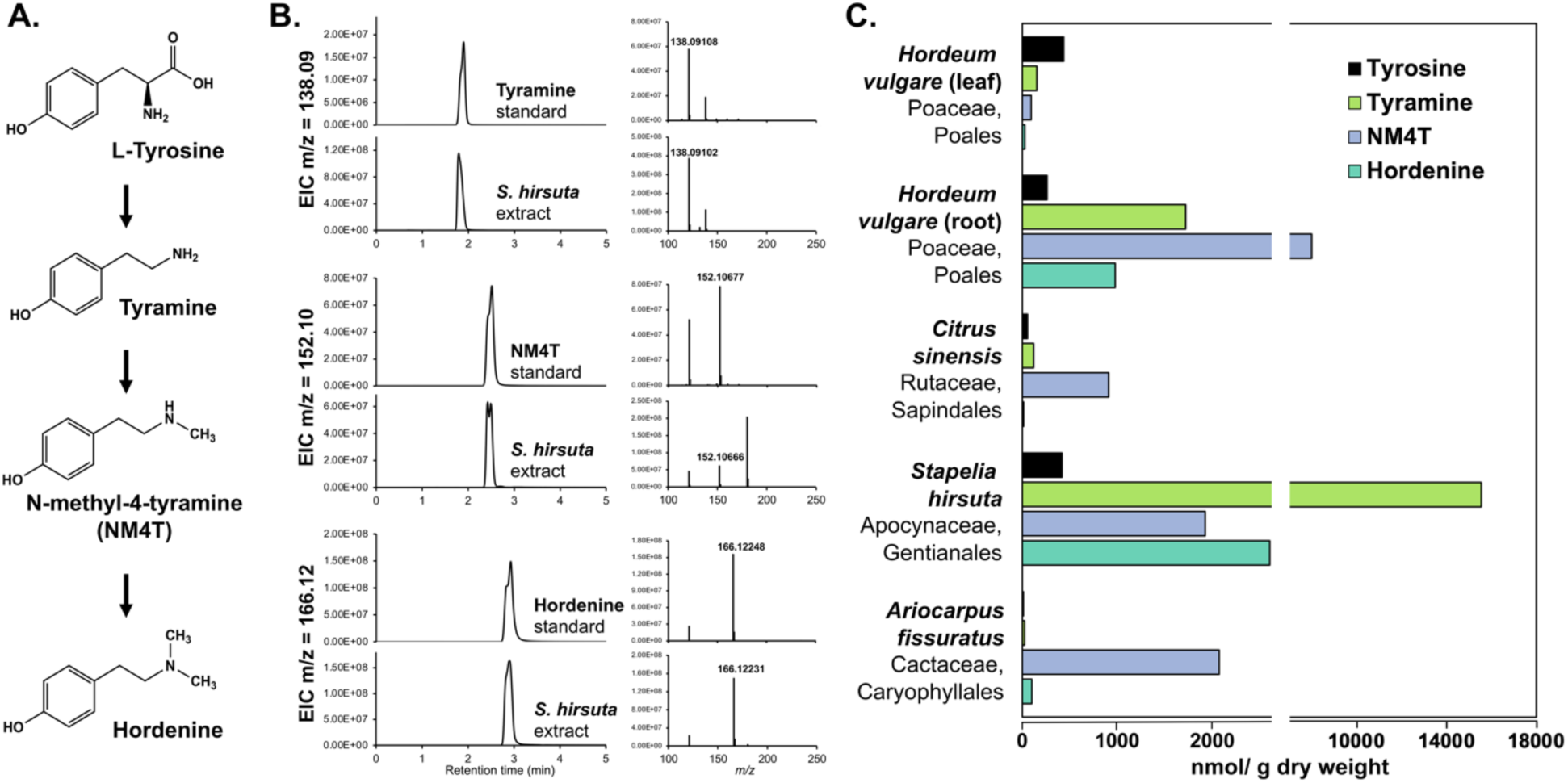
Experimental validation of hordenine and its precursors. **A.** Hordenine is synthesized from the aromatic amino acid L-tyrosine via tyramine and N-methyl-4-tyramine (NM4T) as intermediates. **B.** Metabolites were extracted from the leaf tissue of *Stapelia hirsuta* (Appocynaceae) and analyzed for the presence of hordenine and its upstream precursors using LC-HRMS by comparison with corresponding authentic standards (100 µM concentration). Mass searches were performed using the exact mass of each compound with an allowable error of 10 ppm at 5 decimal places, generating extracted ion current (EIC) traces shown. **C.** Further metabolite analysis of leaf and root tissues of *Hordeum vulgare* and the leaf tissues of other plant species showed that some non-Poaceae plants also accumulate comparable or even greater levels (nmol/g dry weight of plant tissues) of hordenine and its precursors than *H. vulgare* (Poaceae).

Finally, the phylochemical map of tyrosine-derived compounds presented here also revealed high lineage specificity in the dopamine-derived compound class, with clusters in Caryophyllales, Liliales, and Fabales (**Figure 2C**, red arrows). For example, Liliales was reported to accumulate several dopamine-derived compounds, including the intermediates of colchicine biosynthesis, such as (*S*)-Autumnaline and Demecolcine. In addition, 3,4,5-Trimethoxyphenethylamine (mescaline), which is well known to be produced in peyote cactus (*Lophophora spp*.) in Cactaceae (Caryophyllales order),^32,33^ was also detected in Fabales, specifically in the Fabaceae family, the *Acasia* genus within the mimosoid clade.^34,35^ Finally, both Caryophyllales and Fabales were found to produce other dopamine-derived compounds such as anhalonidine and anhalamine (**Figure 2C**, red arrows). These examples illustrate that the pipeline can highlight lineages with concentrated occurrences of a group of related chemicals, a potentially valuable tool in detecting examples of convergent metabolic evolution beyond those already reported.^36^

### The potential of large language models to complement text mining approaches

So far, our work indicated that using text mining to find co-occurrences of the names of plant chemicals and species could generate a large number of candidate compound-species associations (3,631 in a set of tyrosine-derived queries). However, by manually inspecting each full text yielding these candidates, we found that 23.7% of hits were false positive associations, that is, the name of a given compound and species appeared close together in an article’s text, but the article did not provide empirical data for the compound actually being detected in an extract from that species. Still, even with a high false positive rate, the text mining approach yielded useful and sometimes unexpected, yet accurate, information as described above. These outcomes supported the rationale for constructing larger phylochemical maps with a text mining approach but suggested that methods to reduce the false positive rate (other than laborious manual inspection) should be explored. Recent studies in natural language processing have highlighted the abilities of large language models to answer questions about the meaning of text strings, including text describing chemical and biological concepts,^37–39^ and such models are now readily accessible. Accordingly, we next evaluated the abilities of large language models to categorize candidate compound-species associations by using the manually curated dataset as a ground truth set (**Figure 4A**). If effective, such models could greatly enhance our ability to generate accurate, large-scale compound-species association maps directly from existing literature.

**Figure 4:**
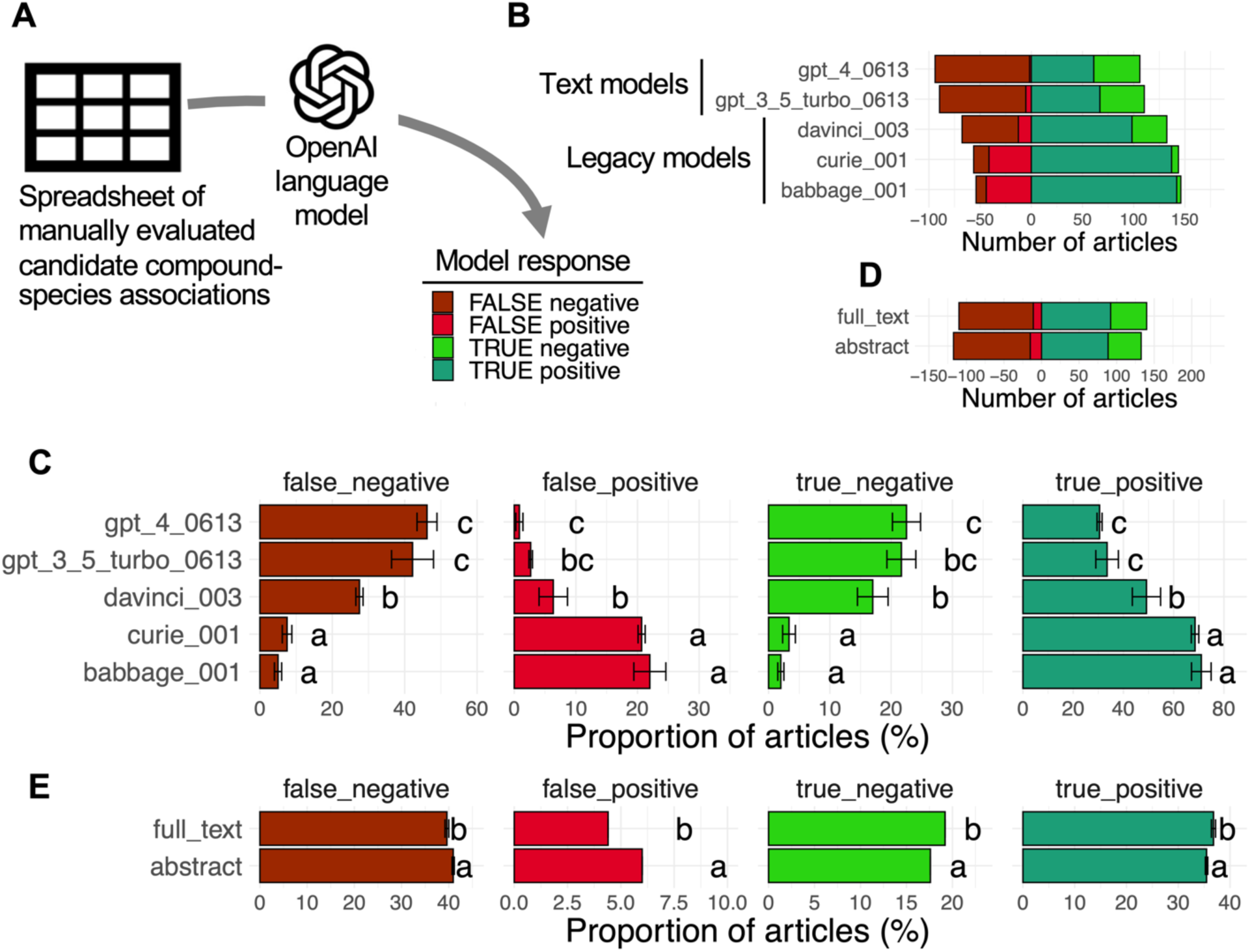
Language model-based evaluation of compound-species associations. **A.** Schematic of language model evaluation process. **B** and **C.** Comparison of OpenAI language models’ ability to assess compound-species associations based on title and abstract text in terms of number of articles (B) and proportion of articles (C). **D** and **E.** Comparison of compound-species association assessment abilities of gpt-3.5-turbo-0613 using only title and abstract text versus when full texts are also supplied in terms of number of articles (D) and proportion of articles (E). Note that the proportion of false positives and true negatives was identical between the independent samples, so the error bar sizes are zero. Bar heights and error bars represent means and standard deviations of n=3 independent tests (‘replicates’) conducted using random sampling, respectively. Letters (‘a’, ‘b’, ‘c’) above bars indicate significant differences as determined by ANOVA and post-hoc Tukey tests, p < 0.05.

Of the large language models that are publicly available, we tested three of OpenAI’s legacy models, babbage, currie, and davinci, as well as two text models, gpt-3.5-turbo-0613 and gpt-4-0613, because these were the most accessible and advanced models at the time of study. To test the models’ abilities, we began by filtering our manually evaluated compounds-species associations for only those 3,275 associations that could be distinguished as true positives (2,409) or false positives (863) from manual inspection, thus eliminating associations whose validity was ambiguous. To test a language model’s abilities, we created a query containing the text of an article’s abstract and title, as well as a question: ‘Here is an article’s title <title> and its abstract <abstract>. Answer yes or no: is the compound <compound name> found in <taxon name>?’. In a first round of queries, we randomly selected 200 putative associations from among the 3,275 candidates and passed each association, as a separate query, to each model. We conducted a total of three rounds of queries using a different random selection each time. These random selections served as independent experiments (or ‘replicates’) in this context. Then, we compared the models’ responses with the results of our manual evaluation to categorize the models’ responses as correct or incorrect (**Figure 4A**).

Our first observation was that, compared with the legacy models, the text models (e.g., gpt-4-0613), yielded a substantially lower number of false positives (a negative association erroneously identified by the model as positive, <5% versus >5%). However, the lower false positive rate was accompanied by lower proportion of true positives (correctly identified positive associations; ∼30% versus > 30%; Tukey HSD test, p < 0.05, Bonferroni correction applied; **Figure 4B and C**). Relative to the legacy models, the text models also produced a higher number of negatives—both true negatives (correctly identified instances in which the co-occurrence of a compound name and a species name did *not* indicate the presence of that compound in that species) and false negatives (erroneous claims that the co-occurrence of a compound name and a species name did *not* indicate the presence of that compound in that species). At the risk of anthropomorphizing, one way to think about the increase in negatives is an increase in ‘skepticism’ as model sophistication increases, or that the more sophisticated models are able to distinguish between textual co-occurrence and biological co-occurrence. Overall, our results indicate that language models, particularly the text models, can greatly reduce the proportion of false positives in a phylochemical mapping dataset in an automated fashion.

In addition to comparing different language models, we examined whether providing the full text of an article improved the ability of a language model to classify candidate compound-species associations. For this test we chose to work with gpt-3.5-turbo-0613 for three reasons: (i) there were no significant differences in the performance of gpt-3.5-turbo-0613 and the most advanced model, gpt-4-0613, in our initial tests (**Figure 4C**), (ii) at the time of our experiments there were rate limits imposed on gpt-4-0613 usage (only 25 queries per hour), and (iii) the availability of gpt-3.5-turbo-16k-0613, which was identical to gpt-3.5-turbo-0613 except that it had a 16,000 token context window that would enable working with full text articles. Using gpt-3.5-turbo-0613/gpt-3.5-turbo-16k-0613, we again performed three rounds of queries. In each round, we randomly selected a set of 250 compound-species associations and constructed two types of queries for each – one that asked whether a given compound was found in the species of interest based on the title and abstract alone, and a second that asked the same question but also provided the full text in the query. When provided with full texts, the model was able to produce a slightly higher proportion of true negatives and true positives, as well as a slightly lower proportion of false negatives and false positives (Tukey HSD test, p < 0.05, Bonferroni correction applied; **Figure 4D and E**). In other words, providing full texts seems to result in a small but detectable improvement in language model-based classification of candidate compound-species associations.

Overall, our work with the language models revealed them to be an excellent complement to text mining in building phylochemical maps. For the scientific community to utilize and further explore this approach, we provide the code we used to perform our language model assessments at https://github.com/thebustalab/phylochemical_mapping. We also provide raw data in the form of candidate compound-species associations with annotations, and true compound-species associations missed by the text mining pipeline, as well as raw results from the language model evaluations and the code used to create the figures presented in this article. These open resources will allow other scientists to further expand the map and associate with other traits as discussed below.

## Discussion

In this study, we developed a proof-of-concept workflow for building maps of chemical occurrence across a plant phylogeny using text mining and downstream processing. By using tyrosine-derived compounds as test cases, we were able to validate that this approach can indeed identify the known presence of compounds in well-studied lineages and highlight previously unrecognized patterns of chemical distribution, illustrating the efficacy and power of our workflow. In addition, the application of large language models, combined with our manually curated dataset, showed that downstream evaluation after text mining is critical but can largely be automated in future iterations. Here, we discuss the limitations of our workflow and potential solutions, as well as future applications and how we can make more comprehensive phylochemical maps that capture larger bodies of literature and plant chemical diversity.

### Limitations of the text mining workflow and potential solutions

Our manual curations (i) showed that the text mining identifies a high rate of false positive hits for compound-species associations (**Supplemental Figure S2**) and (ii) identified several types of false positive associations. Notably, further application of language models to evaluate the potential associations, derived from the text-mining, produced a low rate of false positives, indicating that these models are quite conservative in their assessments and tend to accept very few incorrect compound-species associations. However, this conservative approach also produced a high rate of false negatives, meaning that the models dismissed a considerable number of correct associations. Such filtering resulted in sacrificing the size of the output map for the sake of accuracy, leaving room for improvement in the sophistication of the evaluation portion of our workflow. One route for improvement could be to use techniques like chain of thought prompting^40^, or dedicated techniques for the extraction of structured data from unstructured texts^41^. Alternatively, prior works have shown that performance gains in classification tasks can be achieved by fine tuning a language model using annotated examples42-44. Therefore, the evaluation portion of our future workflow could be enhanced with a similar approach using the several types of false negatives and false positives that we manually annotated in this study to create a fine-tuned classification model.

We also sought to evaluate the comprehensiveness of the text mining process itself by looking for relevant studies that our text mining method might have missed. Through manual searches using Google Scholar for each of the 63 tyrosine-derived compounds (excluding 8 compounds with very high number of medical articles, **Supplemental Figure S1**), we discovered 291 overlooked articles (**Supplemental Table S3**). These articles reported a total of 344 valid compound-species associations, which we again confirmed through manual inspection. However, these articles and associations were not identified through our text mining process (**Supplemental Table S2**), likely because they were not included in the literature database (i.e., SciFinder) that we initially used to search for information on tyrosine-derived compounds. When we mapped these 344 newly found associations, we found similar overall trends compared to our existing text-mined dataset (**Supplemental Figure S3**). Nevertheless, these results suggest that when we scale to cover more species and a broader range of chemical compounds, we need to expand the range of databases that we query to build more comprehensive phylochemical maps. Also, we need to always keep in mind that a lack of reports does not necessarily indicate the absence of a compound in a certain species.

Another limitation of the method described here was that certain compounds had a very high number of papers describing them (e.g., epinephrine, morphine, **Supplemental Figure S1**). In our workflow, we elected not to map these compounds described in more than 10,000 papers, because many of the reports detailed medicinal uses rather than the presence of these compounds in specific plant species. In other words, the signal-to-noise ratio for widely studied medicinal compounds is low. Our decision to omit such compounds led to notable absences in our maps: for example, the absence of positive hits for isoquinoline alkaloids (such as codeine and morphine) in Ranunculales, an order that includes poppy plants and is well-known for producing these compounds.^45^ This limitation could be addressed in future works by developing procedures to effectively map compounds that are the subject of a large volume of research, such as by eliminating only certain articles (e.g., those from medical journals) for these selected compounds or by training language models to identify and dismiss medical articles while preserving others. Alternative strategies could involve only mining certain portions of an article (its results but not introduction section, for example) when generating candidate compound-species associations.

Finally, a lack of sufficient reported data for some compounds may create gaps within the map and their underlying patterns of chemical distribution may be invisible (even if they exist). This situation can be compared to the concept of sequencing depth: without sufficient data, it is challenging to accurately identify genetic features. Furthermore, since phylochemical maps are two-dimensional, there is a parallel issue regarding phylogenetic resolution. In cases where the phylogenetic data is limited (e.g., certain plant families), even abundant chemical data might not reveal patterns of interest. While notable patterns of certain chemicals at the level of plant orders emerged in this study (**Figure 2C**), distributions at the levels of plant families or genera may require further species sampling and refinement in the molecular phylogeny. Therefore, our study also highlights the importance of continued expansion of both axes—chemical and phylogenetic —within the scientific community to achieve adequate depth for generating useful maps.

### Applications of the phylochemical database

One interesting finding from our mapping efforts was the presence of dopamine-derived compounds in Caryophyllales, Liliales, and Fabales (**Figure 2C**). These results raise the possibility that these distant lineages somehow independently acquired highly active upstream pathways to produce elevated amounts or availability of the dopamine precursor. Prior studies showed that Caryophyllales and Fabaceae within Fabales have highly active biosynthetic pathways of tyrosine, an upstream precursor of dopamine.^46–48^ The current finding suggests that Liliales may also have altered regulation of the upstream precursor supply pathway, leading to enriched occurrence of dopamine-derived compounds. This demonstrates one example where phylochemical mapping can provide a novel hypothesis in an unbiased or untargeted manner, which could be further evaluated experimentally in future works. Similarly, once we build a larger, more comprehensive phylochemical database in the future, we can isolate data from certain plant orders or families and assess the characteristics of certain regions of metabolism within that lineage, which will offer a system-level view of the phylochemical map and underlying metabolic evolution.

Another interesting pattern that emerged from our mapping efforts was repeated reports of tyramine, N-methyl-4-tyramine, and hordenine in multiple lineages (e.g., Sapindales, Gentianales, Poales, **Figures 2C and 3C**). While we know that these compounds are the intermediates and product (respectively) of the hordenine biosynthetic pathway (**Figure 3A**), this result illustrates that phylochemical mapping has the potential to illuminate lineage-specific metabolic innovations, such as lineage-specific modification or evolution of a specific branch of a metabolic network. Increased resolution of the mapping, as discussed above, could also predict *when* such evolutionary innovation might have occurred. These results suggest that, for chemicals generated by yet unknown biosynthetic networks, phylochemical maps could identify potential pathway intermediates. Together with a chemical structure logic system, such as SMILES,^49^ we could predict potential structures of unknown or novel biochemical pathways and reconstruct the lineage-specific evolution of hypothetical networks. A phylochemical database could be also linked to metabolic network databases, such as Plant Metabolic Network^50^ to facilitate our understanding of phylogenetic distribution and thereby evolutionary history of plant metabolic networks. Finally, we note that, in addition to metabolic networks, metabolomic data could be integrated with a phylochemical database which could potentially help assign unknown peaks in metabolomic datasets by linking the phylogenetic occurrence of those peaks to previously reported areas of metabolism within a given lineage.

A phylochemical map could be further merged with maps of other traits such as genetic, biochemical, morphological, and physiological traits, as well as climatic and geographic distribution patterns. Comparing the phylogenetic distributions of multiple traits with compound occurrence could potentially illuminate novel and important links. For example, a recent study applied a similar idea to identify a link between certain geographic patterns and enzyme biochemical traits.^17^ Also, since metabolites are synthesized through enzymes encoded in genome of each individual, the phylogenetic distribution of certain chemicals with genomic and biochemical information could help identify the genetic basis of chemical synthesis. Finally, the comparison of chemical distributions and ecological and geographic patterns could infer potential biological functions and ecological roles of plant specialized metabolites.

In addition to providing research-oriented benefits, maps of multiple trait types are also excellent catalysts for interdisciplinary systems thinking, which have the potential to help us strengthen the STEM workforce.^51^ The manual curation process of this study was carried out as a part of virtual undergraduate research during the COVID pandemic, where five students worked closely to critically evaluate scientific literature, contributed to the interpretation of the findings, and also learned various functions and distribution of diverse plant chemicals at a larger phylogenetic context. To further facilitate such research and educational efforts, software tools that allow interaction with maps of multiple data types could be highly useful. For example, a previously published fatty acid database, PlantFADB, illustrates one potential platform by which researchers and students could interact with phylochemical maps.^52^ One excellent feature of such a website is the ability to view the underlying data on user selected phylogenetic level, such as the abundance or presence of a given compound across different plant orders, different plant families, or different genera. Depending on when a biochemical pathway of interest evolved, viewing the data at one phylogenetic level might be more useful than other views. For example, higher-level phylogenetic views might not provide any resolution for more lineage-specific compounds, while a view at the genus or even species-level might only highlight patchiness in the data and such a map might not be a good tool to test a given hypothesis. In short, larger phylochemical databases developed in the future could benefit from the option to be viewed in a dynamic fashion.

### Generating more comprehensive phylochemical maps

While this study built a phylochemical map for a targeted class of compounds (tyrosine-derived compound) as a case study, the workflow we developed is capable of generating plant chemical occurrence maps in a much more comprehensive and untargeted manner. This untargeted approach would involve processing scientific articles with a language model to extract compound-species associations without focusing on specific chemical compounds or plant species. Such an approach would yield untargeted maps, providing a broader view of plant chemical distributions to help us identify novel molecular mechanisms that underpin chemical evolution, predict pathway plasticity, unearth novel pathways,^53^ and identify variant enzymes with potential industrial or agricultural applications.

The creation of large, untargeted plant chemical maps from extensive literature must consider the scalability of the workflow, which comprises three main steps: accessing or compiling a literature collection, extracting candidate compound-species associations, and evaluating these associations. Scaling the literature database(s) that can be used for input into a mapping workflow seems essentially complete with the availability of near-comprehensive and programmatically accessible public databases like PubMed, even if only abstracts and not full texts are available. Scaling up the extraction process of candidate compound-species associations is also practical, since the list of plant chemicals and species can be found in large public repositories, such as PubChem and botanical databases like World Flora Online. Finally, the evaluation of these associations can be done through a computer program capable of understanding natural language, though small-scale manual curations could be helpful for validation and further training of language models. Fortunately, unlike tasks with exponentially increasing solution spaces, such as phylogeny building, the evaluation of compound-species associations based on text excerpts is linear in complexity. Therefore, the required time and effort can be scaled linearly with the number of evaluations, which makes the extensive expansion of phylochemical mapping feasible. In fact, we could theoretically expand the workflow to encompass all known chemical space as well as species outside of the plant kingdom. This scalability highlights the relevance of our work and its potential value to researchers across various fields of natural product research that span the tree of life.

## Materials and Methods

### Text mining for compound-species associations

To prepare a list of tyrosine derived compounds for subsequent literature mining, we inspected prior published literature and the KEGG plant chemical compound list to identify 71 tyrosine derived compounds that spanned a range from very common or ubiquitous compounds produced in nearly all lineages to highly lineage specific compounds known to be produced in only a select few genera. SciFinder searches were performed manually by querying the database for the CAS number associated with each of the 71 tyrosine derived compounds. The results page from each search was filtered using the SciFinder filtering tool to obtain articles associated with the keywords ‘plant’ or ‘taxonomy’. Compounds with more than 10,000 hits were excluded manually using SciFinder. The records for the remaining articles were temporarily downloaded and compared against the World Flora Online Plant List (https://wfoplantlist.org/plant-list) using regular expression-based pattern matching in R^54^. Synonyms and unofficial names were not considered in our pattern matching. The 2,571 articles that mentioned a plant genus and species name present in the World Flora Online Plant List were retained and organized into a spreadsheet (**Supplemental Table S2**), and the other records were deleted from the local storage space.

### Manual curation of compound-species associations

Full texts for each of the 2,571 articles were obtained through our university libraries. Each was manually inspected to determine whether the putative compound-species associations found by the pattern matching approach were true, false, or unclear. Additional metadata was also collected from each full text, including data on the tissue types reported to accumulate the compound of interest, other plant species reported to accumulate the target compound within each article, and whether the report of the compound included analytical quantification. To visualize the resulting data, we pruned a published phylogenetic tree to include only the orders that were represented in our association data. Phylochemical maps were visualized by using the R package ‘ggtree’^55^ to visualize a pruned phylogeny^26^ plotted alongside a heatmap representing the association data generated with ggplot2^56^.

### Analysis of metabolites by UHPLC-MS

Approximately 100 mg of healthy fresh plant tissue was harvested from plants grown in the Botany Greenhouses of University of Wisconsin-Madison and flash frozen in liquid nitrogen. The tissues were then freeze dried using the Benchtop Freeze Dryer (Labconco) for no less than 48 hours and pulverized with a bead beater. Ten to twenty milligrams of frozen powders were aliquoted to new frozen tubes, weighed, and resuspended in 800 μL of LC-MS grade methanol: chloroform, (v/v 2:1) with 10 μM ^13^C(6)-tyrosine (Cambridge Isotope, CLM-1542-0.25) as an internal recovery standard. After rigorous vortexting for 10 minutes, the extracts were centrifuged for 10 minutes at 21,100 *g* at room temperature, and 800 μL of the supernatant was transferred to a new tube, mixed with 600 μL water and 250 μL chloroform, vortexed for 30 seconds, and then centrifuged again for 5 minutes at 21,000 *g* to generate phase separation. The upper aqueous phase was transferred into a new tube and dried down in a CentriVap SpeedVac (Labconco) overnight at room temperature. The dried samples were resuspended in 70 μL of 80% methanol by vortexing for 30 seconds and sonicating for 5 minutes, followed by centrifugation for 5 minutes at 21,000 *g*. 30 μL of the supernatant was then transferred to glass vials for injection onto a Vanquish Horizon UHPLC Binary Pump H coupled to a Q-Exactive mass spectrometer (Thermo Scientific). All reagents were of LC-MS quality. One microliter of the sample was injected onto a HSS T3 C18 reversed phase column (100 × 2.1 mm i.d., 1.8-μm particle size; Waters) and eluted using a 26-minute gradient comprising 0.1 % formic acid in water (solvent A) and 0.1 % formic acid in acetonitrile (solvent B) at a flow rate of 0.40 mL/min and column temperature of 40 °C, with the following linear gradient of solvent B: 0-1 min, 1 %; 1-10 min, 1-10 %; 10-13 min, 10-25 %; 13-18 min, 25-99 %; 18-22 min, 99 %; 22-22 min, 99-1 %; 22-26 min, 1 %. Full MS spectra were recorded between 0.55 and 18 min using the full scan positive mode, with the following parameters: sheath gas flow rate, 45; auxiliary gas flow rate, 13; sweep gas flow rate, 1; spray voltage, 3.75 kV; capillary temperature, 350°C; S-lens RF level, 50; resolution, 140,000; AGC target 1 × 10e6, maximum scan time 200 ms; scan range 100–1000 m/z. Metabolites were identified through a mass search of the compounds’ mass to charge ratios (m/z), generating an extracted ion chromatogram (EIC) from which retention time and mass spectra were matched to high purity authentic standards: tyrosine (A11141, Alfa Aesar), tyramine (AC140610050, Acros Organics), and n-methyl-4-tyramine (OR-4655, Combi-Blocks), hordenine (04476, Sigma Aldrich). Quantification of each compound was done by manual integration of peak area using Xcalibur 3.0 (Qual Browser, Thermo Scientific) and abundance was calculated by reference to the authentic standards. The final compound contents were determined by normalizing based on the internal standard (% recovery) and tissue weight for each sample.

### Language model evaluation of candidate compound-species associations

Candidate compound-species associations were evaluated using language models created by OpenAI^57^. Compound species associations were evaluated inside a loop in which each candidate association was passed to the OpenAI API using an R script like:

~~~
query <- paste0(
  “Here is the title: ”, data$title[i],
  “Here is the abstract: ”, data$abstract[i],
  “ classify your response as \“yes\” or \“no\”: was ”,
  data$compound_name[i], “ found in ”, data$species[i],
  “? Do not include a period in your response. Respond only with yes or no and no other explanation or text.”
)
~~~

The system prompt used was “You are an expert in scientific literature and can easily and accurately parse text for scientific literature tasks.” The output from the model was used directly in downstream analysis. A full example of the code used to evaluate the language models is included at the project GitHub site. Once the responses from the language model had been tabulated, ANOVA and post-hoc Tukey tests, implemented in R via the ‘statix’^58^ were used to test for any significant differences.

## Data Availability Statement

The datasets generated and analyzed during this study as well as the relevant code are available on GitHub at https://github.com/thebustalab/phylochemical_mapping. This includes the annotated 3000+ compound-species associations, code used to evaluate language models, and code used to create phylochemical maps presented in this work.

## Author Contributions

L.B. and H.A.M. conceived the research. L.B. conducted the text mining and phylogenetic mapping with the help of H.A.M. The curation of candidate compound-species associations was performed by D.H., C.M.H., A.G., Y.W., M.G., and M.S. Language model evaluation was conducted by B.J. and L.B. LC-MS validation of hordenine and its precursors was done by M.S. with an initial assistant from M.G. The manuscript was drafted by L.B., H.A.M., D.H., and M.S. with input from all authors. All authors contributed to the analyses and interpretation of the data and read the final version of the manuscript.

## Acknowledgements

We thank Drs. Ricardo Kriebel and Ken Sytsma for helpful discussions for mapping chemical traits across the plant phylogeny and Dr. Bethany Moore for discussing and testing a possibility of a machine learning approach. We are also grateful to Dr. Alan Oyler for assistance with preparing this manuscript. L.B. gratefully acknowledges support from the Swenson College of Science and Engineering in the form of start-up funds and a SciFinder Future Leader Award from CAS SciFinder. B. J. acknowledges the support of a UROP award from the University of Minnesota. This work was supported by grants from the US National Science Foundation (IOS-PGRP-1836824 and DEB-1938597) to H.A.M. The authors declare no conflict of interest.

## Supplemental Material

**Supplemental Table S1.** Compounds considered for phylochemical mapping.

**Supplemental Table S2.** Compound-species associations examined in this study.

**Supplemental Table S3.** True compound-species associations missed by text mining that were identified by manual searching.

**Supplemental Figure S1.**
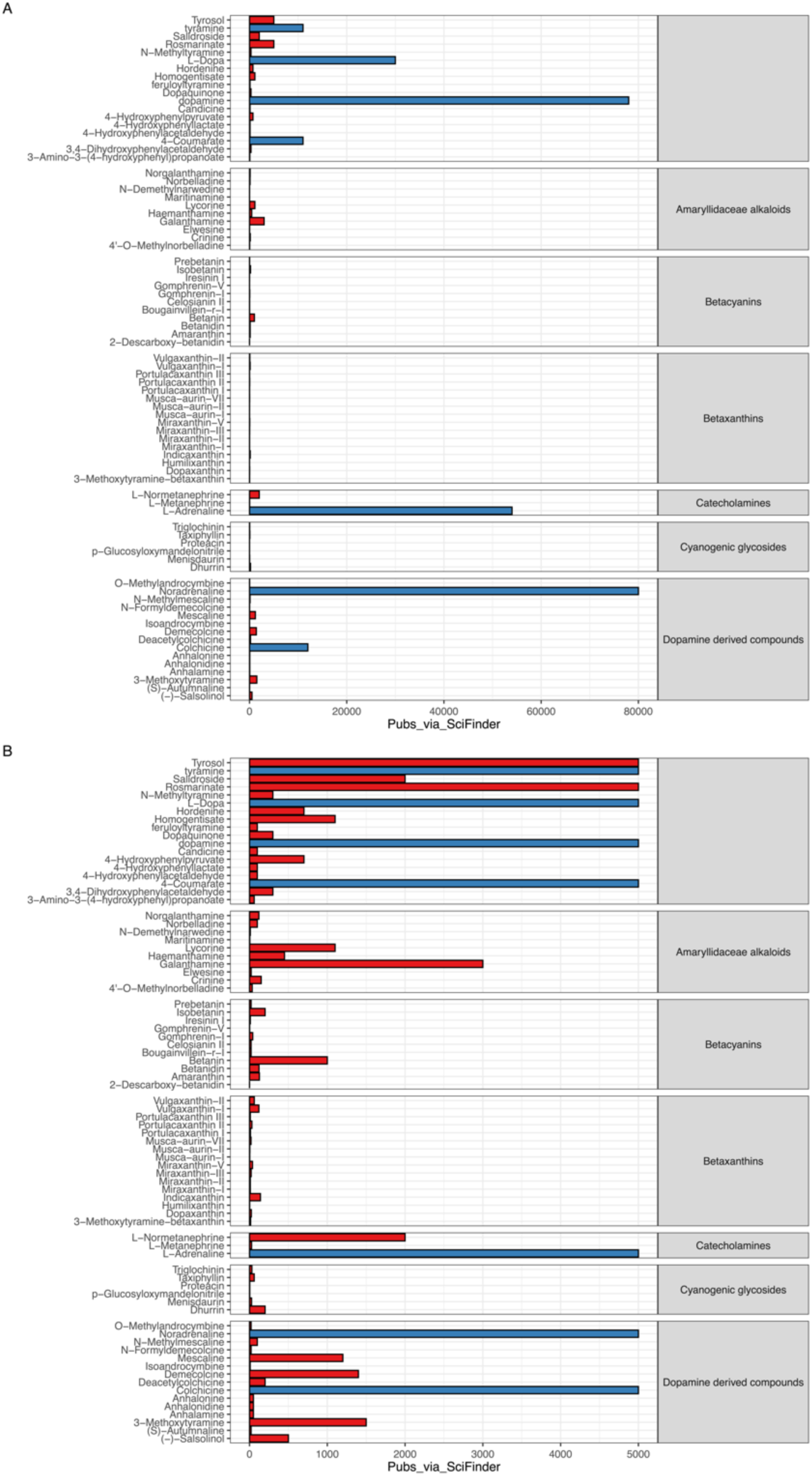
Compounds considered for phylochemical mapping. The two panels present the same data but with different limits on their y scales. Bar heights indicate the approximate number of references for each compound in SciFinder. All titles and abstracts were collected for compounds with fewer than 5000 references. Compounds with more than 5000 references were omitted from subsequent study. Some compounds had only references that did not relate to biological study and were also not considered further.

**Supplemental Figure S2.**
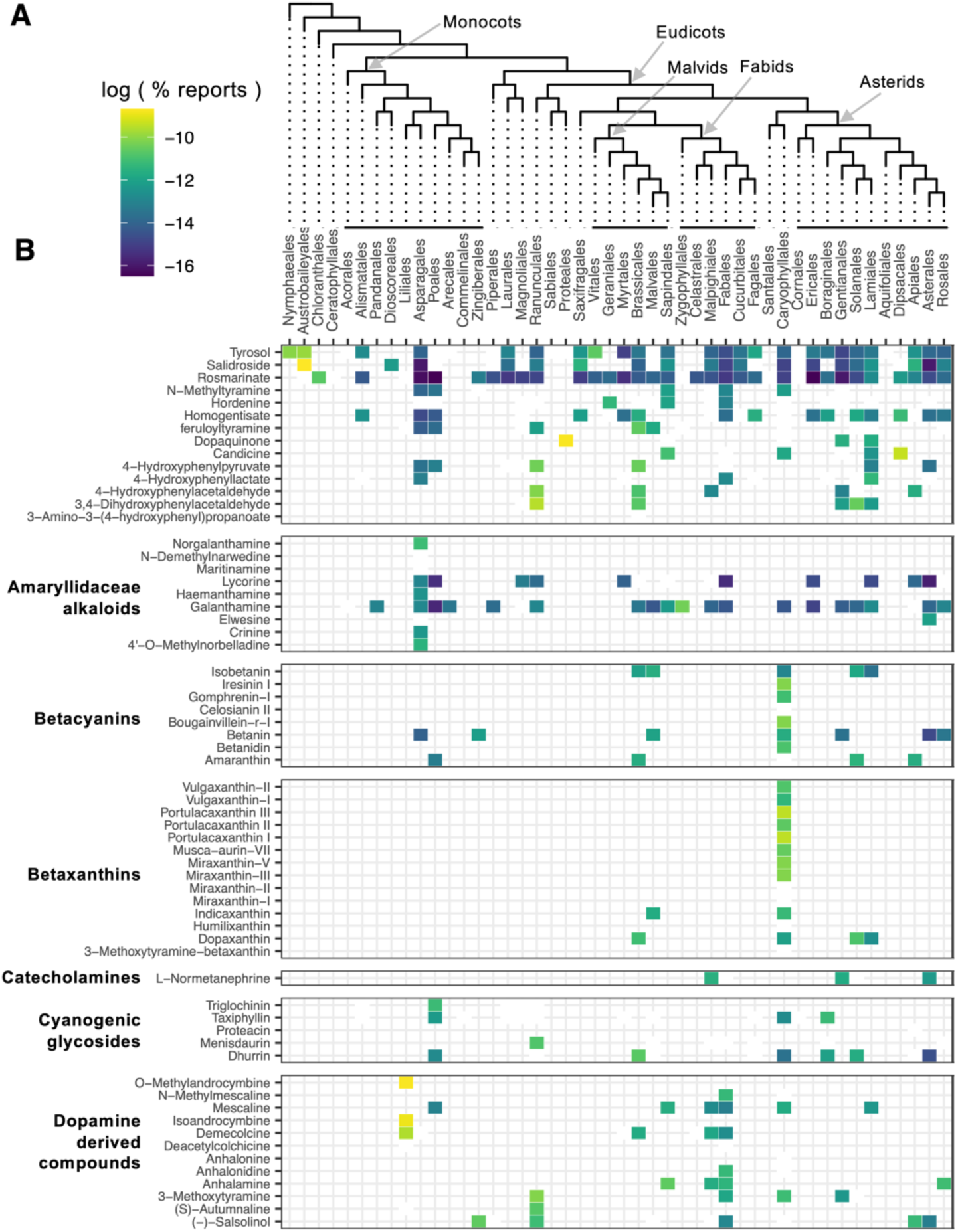
A map of false positive hits from text mining. **A.** Cladogram of angiosperm orders. We mapped compound-species associations onto an angiosperm order phylogeny,^26^ removing branches where no compounds were detected for better clarity. **B.** All false positive compound-species associations. To adjust for differences in species count per order and the number of articles for each compound, we used a normalized metric to color the heat map. This metric is the log of the number of reports for a compound in each order, divided by the total reports of that compound and by the species count in the order, thus emphasizing the compound’s specificity to certain lineages.

**Supplemental Figure S3.**
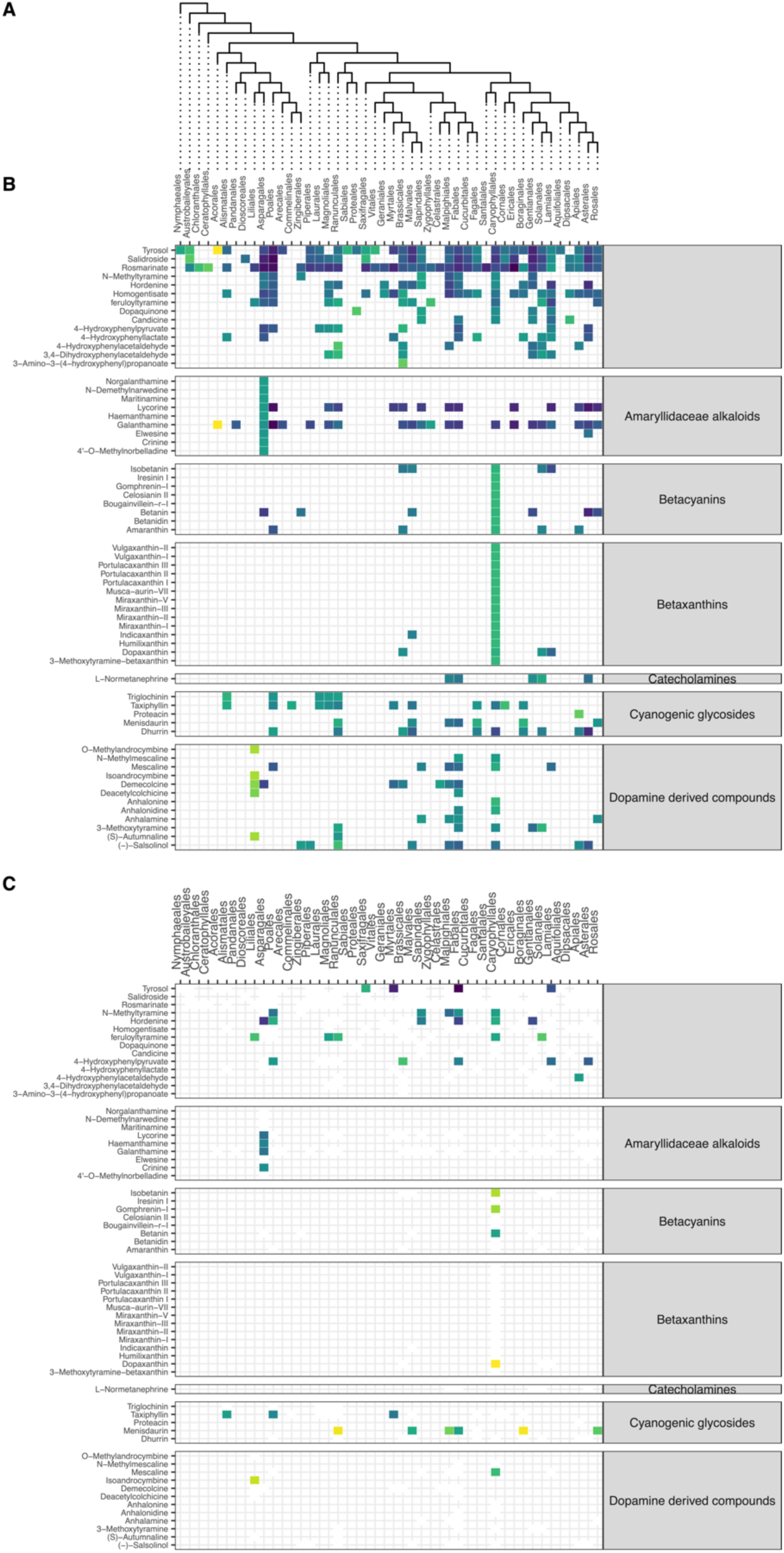
The map of false negatives, true compound-species associations missed by the text mining pipeline. **A.** Cladogram of angiosperm orders. We mapped compound-species associations onto an angiosperm order phylogeny,^26^ removing branches where no compounds were detected for better clarity. **B.** All putative compound-species associations (the same map as shown in Figure 2B as a comparison). **C.** Associations missed by text mining. In B and C, to adjust for differences in species count per order and the number of articles for each compound, we used a normalized metric to color the heat map. This metric is the log of the number of reports for a compound in each order, divided by the total reports of that compound and by the species count in the order, thus emphasizing the compound’s specificity to certain lineages.

